# Fast and haplotype-aware assembly of high-fidelity reads based on MSR sketching: the Alice assembler

**DOI:** 10.1101/2025.09.29.679204

**Authors:** Roland Faure, Baptiste Hilaire, Jean-François Flot, Dominique Lavenier

## Abstract

**Background:** Long-read metagenomic assembly is becoming a critical bottleneck in microbiome analysis, as deep sequencing generates massive datasets that existing methods struggle to assemble while maintaining strain resolution.

**Results:** We present Alice, a lightweight long-read assembler that achieves orders-of-magnitude speedups through a new sequence sketching technique, MSR sketching, compatible with classical assembly methods. Alice assembles a 235 Gbp soil metagenome in 5 hours using only 84 GB RAM—a task that causes most competing methods to exhaust our computational resources (500 GB RAM and 7 days runtime). Across diverse benchmarks, Alice delivered strain-resolved assemblies an order of magnitude faster than state-of-the-art approaches, while producing the most complete assemblies in some cases.

**Conclusions:** MSR sketching overcomes computational barriers in metagenomic assembly, enabling fast, memory-efficient strain-resolved analysis of massive datasets. While Alice’s assemblies were more fragmented than with other assemblers, this approach establishes a promising paradigm for scalable metagenomic analysis.

## 1 Background

With the rise of high-throughput sequencing, genomic experiments have been producing vast amounts of data, far outpacing the growth of computing power predicted by Moore’s law [12]. It is now common for a single experiment to generate dozens or even hundreds of gigabases of data. In parallel, the length and quality of the sequencing reads have improved immensely. PacBio HiFi consensus reads are several thousands of basepairs long with an error rate lower than 0.1% [36]. Oxford Nanopore Technologies (ONT) are also improving in accuracy, down to 1% error [40], and even longer.

Assembling metagenomic datasets, i.e. aligning and merging reads to obtain consensus sequences representative of the metagenome, is a taxing computational task. It can easily require several weeks of CPU hours and hundreds of gigabytes of RAM [13, 18, 39]. As datasets become larger and cheaper to produce, metagenome assembly can become a bottleneck in terms of cost, computation time and quality, a problem made worse by recent sharp increases in RAM prices [25].

A general popular technique to diminish the size of the computations is to sketch the input data, i.e. reduce it to a smaller representation on which computations can still be made [29]. In the realm of genome assembly, sketching has long been employed for all-versus-all read mapping as a first step of the Overlap-Layout-Consensus assembly paradigm [19]. However, it has only recently been effectively integrated into the faster De Bruijn Graph assemblers, specifically for high-fidelity reads. Building on concepts from wtdbg2 (a.k.a. RedBeans) [30], Shasta [31], and Peregrine [9], Ekim, Berger and Chikhi introduced a method that samples a fraction *δ* of the *k*′-mers in the reads, chains the resulting series of *k k*′-mers into “*k*-mers of *k*′-mers” called *k*-min-mers, assemble those *k*-min-mers and subsequently transforms the resulting chain back into a genome sequence [11]. This approach demonstrated remarkable efficiency in a proof-of-concept assembler called mDBG [11], enabling human genome assemblies to be completed in minutes on a personal computer. It was further developed as a metagenomic assembler named metaMDBG [3]. However, these assemblers encounter significant limitations that arise directly from the chosen sketching method.

Metagenomic samples as well as diploid (or polyploid) genome sequences often contain strains that are genetically similar yet functionally distinct [35]. However, when (meta)mDBG sketches the reads as a chain of k-mers, differences —such as single nucleotide polymorphisms (SNPs)—between highly similar sequences is often lost. As a result, both mDBG and metaMDBG do not differentiate highly similar haplotypes.

In this study, we present a novel assembler named Alice. Conceptually, Alice shares similarities with metaMDBG, as it begins by sketching reads and assembling the sketches before decompressing the obtained sequences to yield the final assembly. However, Alice introduces a significant innovation through a new sketching method called Mapping-friendly Sequence Reduction (MSR). Originally proposed to improve read mapping quality [4], the potential of MSR as a sketching technique had not been previously investigated. In our methodology, we employ a carefully parametrized MSR to sketch PacBio HiFi reads, resulting in a computationally efficient assembler that maintains the ability to reconstruct highly similar sequences. The name “Alice” is inspired by Lewis Carroll’s *Alice in Wonderland* [6], where Alice uses a “drink-me potion” to pass through a small door and a “eat-me” cake to return to her original size. In this analogy, Alice represents the reads, the small door symbolizes the constraints of hardware and software capacity, and the potion corresponds to the MSR sketching technique. The assembly process is depicted in Figure 1.

**Figure 1:**
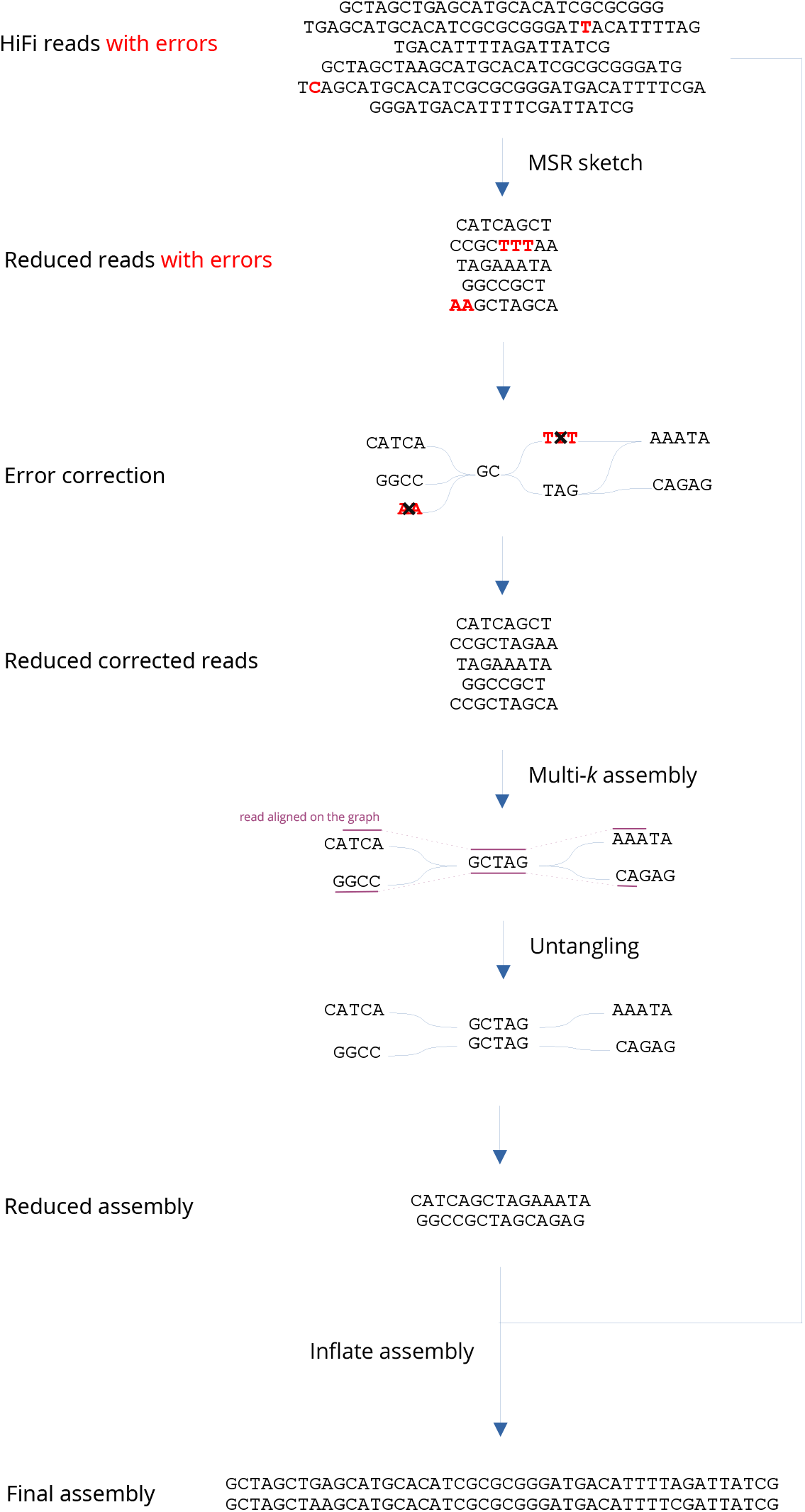
Assembly process of Alice.

### Contribution

We evaluated Alice on three distinct PacBio HiFi metagenomic datasets—(i) a mock community comprising five *Escherichia coli* strains, (ii) a human-gut stool sample, and (iii) a soil sample. Compared with leading assemblers such as metaMDBG [3], metaFlye [18], and hifiasm meta, [13] and myloasm [32], Alice assembled the data one order of magnitude faster and with lower memory consumption. In particular, hifiasm meta, myloasm and metaFlye failed to assemble the soil sample within our hardware limitations (500GB RAM, 7 days runtime). Moreover, Alice reliably discriminated closely related strains and produced the most complete assemblies in some scenarios.

We also examined the assembly of a genomic dataset obtained from HiFi sequencing of the bdelloid rotifer *Adineta vaga*, a rising model organism for which several genome assemblies of increasing accuracy have been published [14, 33], albeit none based on PacBio HiFi yet. On this novel dataset, Alice generated an assembly comparable to those produced by state-of-the-art assemblers LJA [1], Flye [17] and hifiasm [7], but with RAM usage and run time both one order of magnitude lower than these other tools.

## 2 Results

### 2.1 Benchmarking setup

We conducted a benchmarking analysis of Alice, comparing its performance against the four most commonly utilized HiFi metagenomic assemblers, namely metaFlye [18], hifiasm meta [13], metaMDBG [3] and myloasm [32] and four genomic assemblers, namely hifiasm [7], Flye [17], LJA [1] and rust-mdbg [11]. Each assembler was executed using their recommended settings. For Alice, we used the default parameters of a compression factor of 20 and an order of 101, adding the option --single-genome for the assembly of *Adineta vaga*. A detailed discussion of these parameter choices, along with tests of alternative settings, can be found in section 5.6 of the methods.

To benchmark Alice, we first utilized the ZymoBIOMICS Gut Microbiome Standard, a commercially available mixture of 19 bacterial strains and two yeast strains, specifically formulated to mimic the gut microbiome’s composition. PacBio HiFi sequencing data for this standard are publicly available under the accession number SRR13128013. The relative abundances of each organism in the mixture, along with their genomic sequences, are known. Notably, this dataset includes five closely similar strains of *Escherichia coli*, which present a challenge to assemble separately. Secondly, we assessed Alice on two true metagenomic communities: a HiFi sequencing dataset derived from a human stool sample [28] and a HiFi sequencing dataset derived from a soil sample [3]. Both datasets had previously been employed by the authors of metaMDBG to benchmark their own assembler [3]. Finally, as an exploratory endeavor, we also assembled the genome of the animal *Adineta vaga* to see if Alice can be advantageously applied to genomic assemblies.

Assemblies were conducted on AMD EPYC 7552 nodes (500 GB RAM, 7-day runtime limit) using 8 threads (or 16 threads for the soil metagenome assembly).

### 2.2 Alice exhibits an order-of-magnitude speedup over competing approaches

On the soil dataset, myloasm and hifiasm meta exceeded our RAM limit (500GB), while metaFlye exceeded our time limit (7 days). As a consequence, we ran these assemblers on a downsampled version of the original 235Gbp dataset, i.e., 92Gbp for hifiasm meta and metaFlye and 52Gbp for myloasm (as it also failed to assemble the 92Gbp dataset).

Figure 2 presents the runtime and memory consumption of each assembler across the four benchmark datasets. Alice consistently outperformed all competitors, achieving dramatic reduction in both metrics.

**Figure 2:**
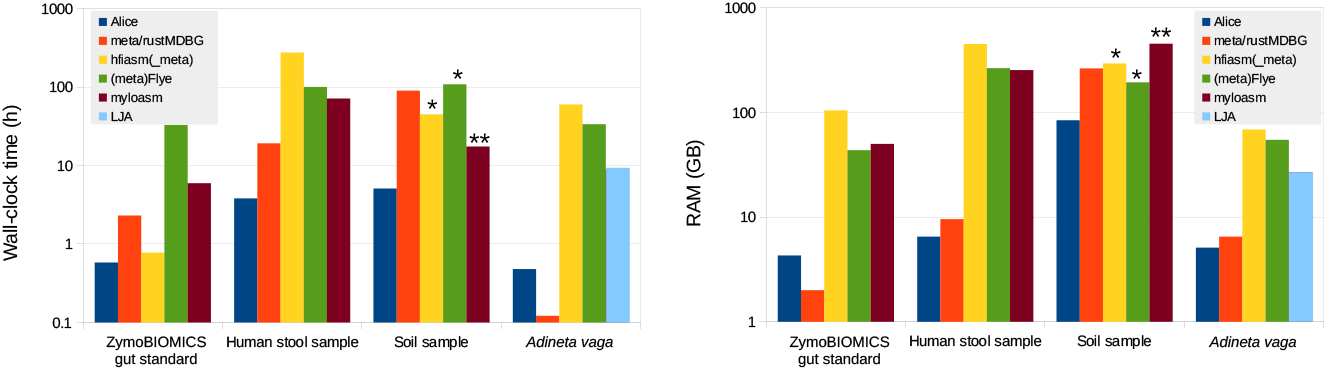
Wall-clock time on 8 threads and RAM usage of the assemblers on the four assemblies assemblies. * and ** denote that the dataset was downsampled to 92Gbp and 52Gbp, respectively, to enable assembly completion. Scales are logarithmic

On the human-stool and soil samples, Alice achieved a speedup of an order of magnitude over metaMDBG, metaFlye, hifiasm meta, and myloasm (5–72×). For the *Adineta vaga* dataset, the speedup was even more pronounced (19–125×), with the exception of rust-mdbg, which produces assemblies of markedly lower quality (discussed below).

Alice’s peak memory usage was 84 GB (soil assembly), requiring three times less memory than metaMDBG, the closest alternative. For all other datasets, Alice and metaMDBG used *≤*10 GB RAM, to compare to the several hundred gigabytes required by metaFlye, hifiasm meta, and myloasm on the human gut dataset.

### 2.3 Alice produces highly complete metagenomic assemblies

We employed metaQUAST [23] to evaluate the assemblies derived from the Zy-moBIOMICS gut microbiome standard. Comprehensive metrics on completeness, duplication ratios, and continuity are reported in Supplementary Tables 1 and 2, with the full metaQUAST output provided in the supplementary data set.

**Table 1:** Contiguities of the stool and soil metagenome assemblies.

|  |  | Alice | metaFlye | hifiasm_meta | metaMDBG | myloasm |
| --- | --- | --- | --- | --- | --- | --- |
| Stool | N50 (kb) | 87 | 122 | 143 | 210 | **402** |
|  | N90 (kb) | 15 | 20 | 14 | 14 | **23** |
| Soil | N50 (kb) | 7.4 | 29 | **41** | 17 | 40 |
|  | N90 (kb) | 3.0 | 13 | **24** | 8.6 | 17 |

**Table 2:** Comparison of assembly statistics of *Adineta vaga* across different tools. 31-mer completeness was computed using KAT. BUSCO completeness was computed against the metazoa odb10 database.

|  | Alice | Flye | hifiasm | LJA | rust-mdbg |
| --- | --- | --- | --- | --- | --- |
| 31-mer completeness (%) | 99.91 | 99.93 | **99.97** | 99.91 | 95.02 |
| BUSCO completeness (%) | 79.0 | 79.1 | **79.9** | 79.0 | 78.3 |
| N50 (Mb) | 1.37 | 9.35 | **11.01** | 0.08 | 0.05 |
| N90 (Mb) | 0.20 | **0.89** | 0.05 | 0.03 | 0.01 |
| Number of contigs | 1436 | **165** | 1429 | 4810 | 11703 |
| CPU time (h) | 1.4 | 253 | 465 | 62 | **0.4** |
| Peak RAM (GB) | **4.8** | 55 | 27 | 26 | 6.5 |

Alice achieved assembly quality comparable to state-of-the-art methods regarding completeness and continuity. Notably, Alice successfully assembled all five conspecific *E. coli* strains separately, each with over 96% completeness. In contrast, metaMDBG, the other lightweight sketching-based assembler in our benchmark, failed to distinguish between strains, resulting in assemblies with completeness as low as 44%.

Evaluating the assemblies of the human gut and soil metagenomes presented more challenge because the exact composition of genomes in those samples was unknown. To assess assembly completeness, we compared the 31-mer content of each assembly with the 31-mers present in the HiFi reads, which were counted using KMC [16]. We assumed that any 31-mer appearing more than five times in the HiFi reads was unlikely to be a sequencing error. Accordingly, we plotted the fraction of these high-confidence 31-mers recovered by each assembler as a function of their abundance in the reads (Figure 3a for the human stool sample and Figure 3b for the soil sample). The contiguities of the assemblies are shown in Figure 1.

**Figure 3:**
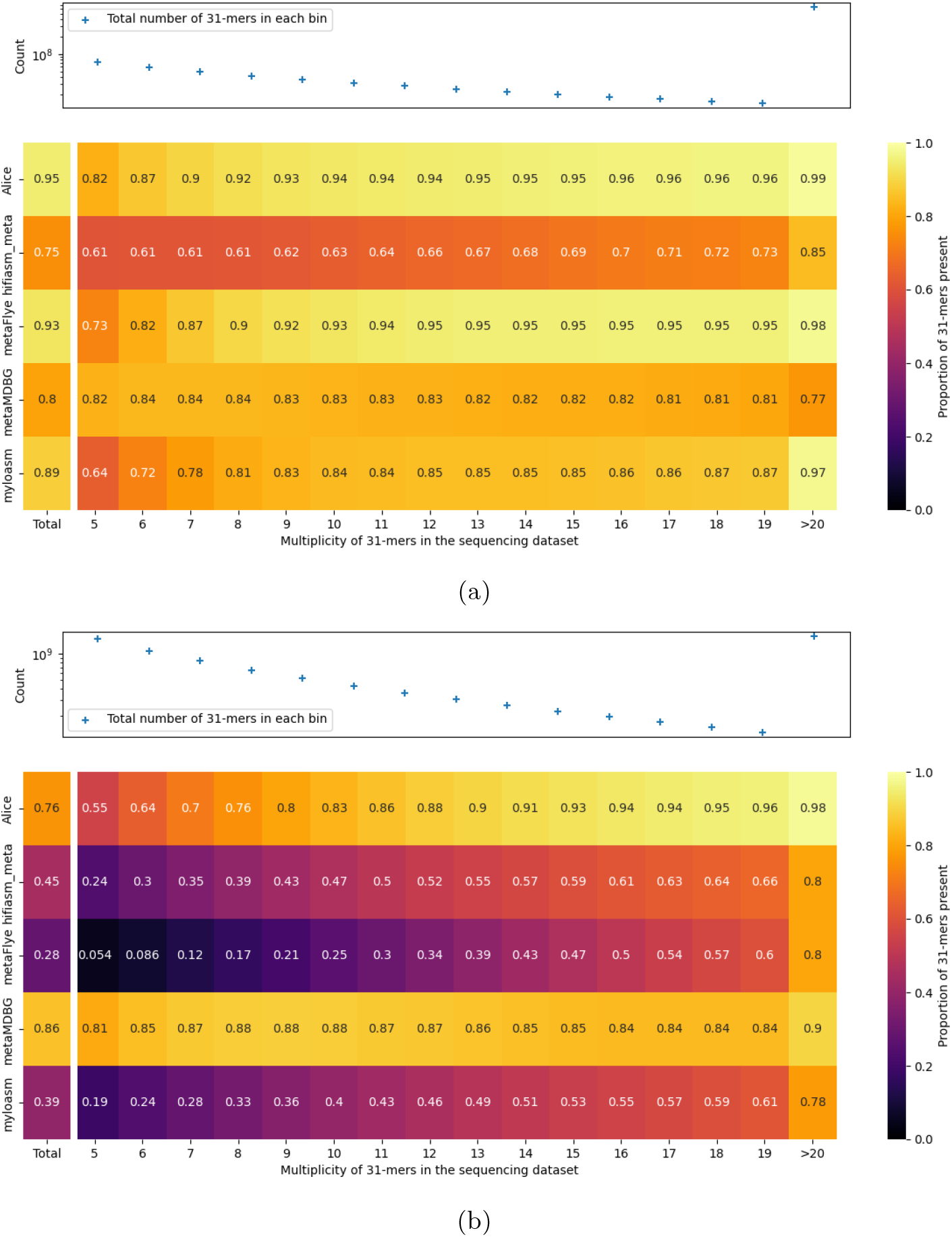
Analysis of the 31-mer abundances of the reads vs the assemblies in (a) in the human stool sample and (b) the soil sample. The top panel displays the number of 31-mers in the reads as a function of their multiplicity in the read dataset. The bottom panel presents a heatmap in which a number of *x*% in bin B indicates that *x*% of the 31-mers of multiplicity B in the reads are present in the corresponding assembly. For example, for the stool sample, 76% of the 31-mers seen 7 times in the reads are found in the metaMDBG assembly. Note that on the soil assembly, only Alice and metaMDBG were able to assemble the full read dataset, other assemblies were computed from a reduced datasets.

Alice produced highly complete assemblies, particularly at high coverage. In contrast, metaMDBG exhibited significant deficiencies at high coverage (≥20×), failing to recover 24% of the 31-mers in the human gut metagenome assembly and 10% in the soil assemblies. These missing sequences were frequently recovered in Alice assemblies as small bubbles, indicating they likely represented strain variants that metaMDBG discarded during assembly.

However, Alice struggled to assemble low-coverage strains, missing up to 45% of 31-mers present only 5 times in reads from the soil dataset. Alice also achieved lower continuity than competing methods, as detailed in Table 1. For example, the N50 of Alice’s soil assembly was only 7.4kb, while hifiasm meta’s was 41kb.

### 2.4 Alice yields highly complete genome assemblies

To benchmark Alice on a single-genome dataset, we sequenced the non-model diploid bdelloid rotifer *Adineta vaga* using PacBio HiFi chemistry to a depth of 140× (the reads are publicly available via BioProject PRJNA1335825). The principal challenge of this assembly is the organism’s relatively high heterozygosity (1.7% [33]), whereas most assemblers are tuned for the far less heterozygous human genome.

We assessed the quality of the *A. vaga* assemblies in two ways. First, we ran a BUSCO evaluation [34, 21] against the metazoa odb10 reference set.

This analysis showed no substantial differences in gene completeness between the assemblers. Second, we performed a spectral analysis of 31-mer frequencies using KAT [22]; the results are displayed in Figure 4. The assembly statistics are summarized in Table 2.

**Figure 4:**
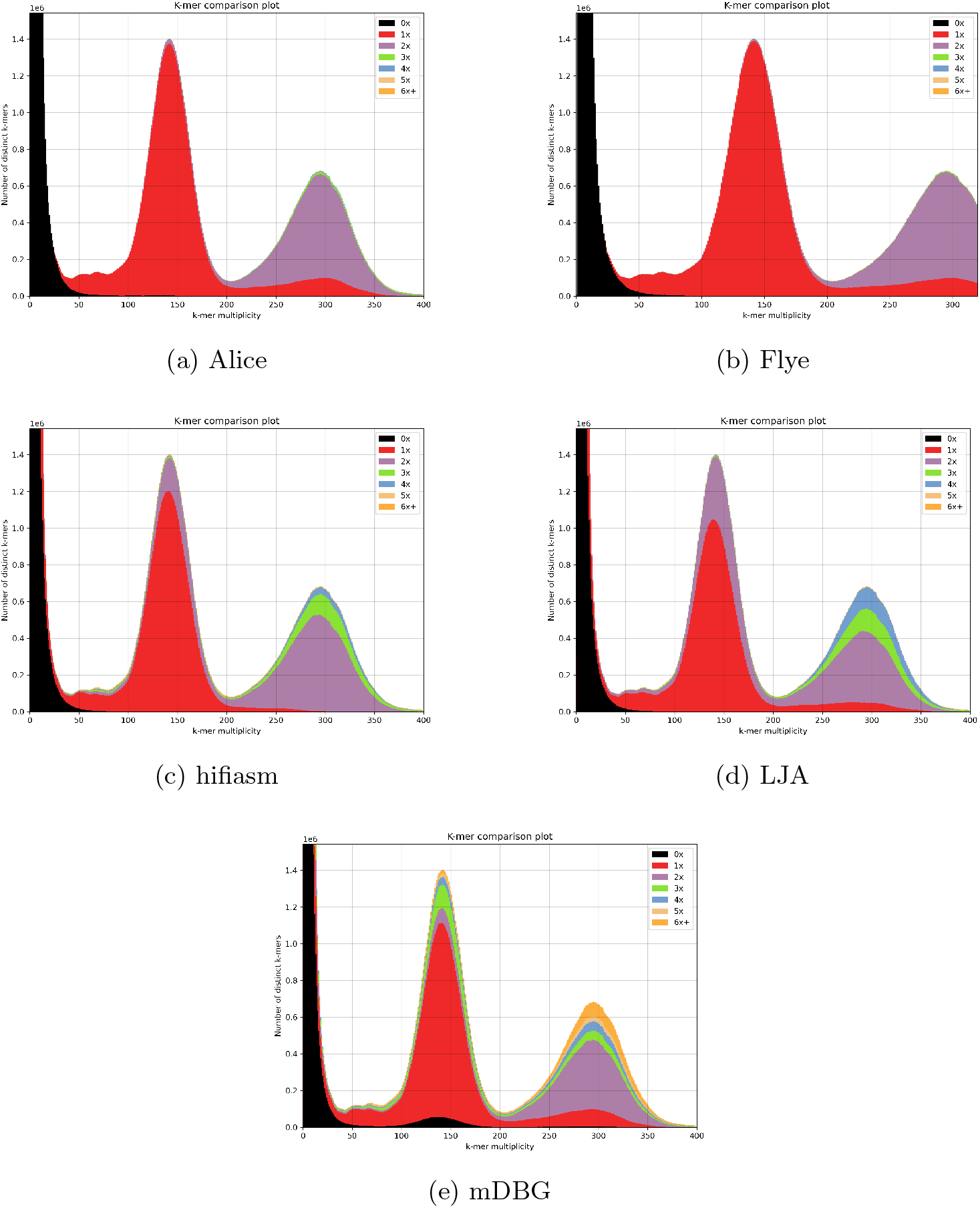
KAT plots of the *Adineta vaga* assemblies against the HiFi sequencing reads. The 31-mer spectra show two multiplicity peaks, one corresponding to homozygous 31-mer (seen twice in the genome and on overage 280 times in the reads), and the other corresponding to heterozygous 31-mers (seen once in the genome and on average 140 times in the reads). Additionally, the peak of 31-mers with a very low multiplicity in the reads corresponds to sequencing errors. The colors show the abundance of the 31-mers in the assemblies.

Our results indicate that Alice produced a genome assembly comparable in quality to the outputs of current state-of-the-art tools such as Flye, LJA, and hifiasm, while requiring substantially less computational effort. Compared with rust-mdbg, Alice achieved similar resource usage but delivered a markedly better assembly. Spectral analysis revealed that even with a high 140× HiFi coverage, rust-mdbg lost a large fraction of 31-mers in its final contigs. Moreover, rust-mdbg, hifiasm, and LJA exhibited pronounced 31-mer over-duplication, whereas both Alice and Flye generated a high-fidelity assembly with only a modest amount of collapsed homozygous regions creating bubble structures in the assembly graphs. Among the tested assemblers, Flye attained the highest continuity, giving it a slight edge over Alice, but at a much higher computational cost.

## 3 Discussion

Alice’s strengths stems from the MSR sketching method. By compressing reads to MSR sketches, it achieves dramatic computational gains—reducing memory use and runtime by orders of magnitude compared to non-sketched-based methods. In contrast, assemblers such as Flye, hifiasm meta and myloasm approached and exceeded their limits in RAM consumption and runtime for the large datasets in our benchmark. metaMDBG, the only competing method capable of assembling the soil dataset, employs k-min-mer sketching, which inherently collapses small sequence differences between closely related strains. This limitation explains metaMDBG’s failure to distinguish the five conspecific strains in the ZymoBIOMICS mock metagenome, whereas MSR sketches preserve these fine-grained distinctions. Thus, Alice combines the computational efficiency of sketch-based assembly with the strain-resolution accuracy of full-read assembly.

Beyond saving time, money, and hardware, the modest resource demands of Alice will enable much deeper sequencing of metagenomic communities in the future. Historically, the main bottleneck for deep metagenomic studies has been obtaining high-quality, high-coverage data, but recent advances now make it possible to generate HiFi datasets of hundreds of gigabases, for a rapidly decreasing price. This increased depth will allow us to detect and characterize low-abundance strains that were previously missed.

Alice’s primary limitation is the limited continuity of its assemblies. This stems from the underlying method for assembling the MSR sketches, a rudimentary de Bruijn graph-based approach that could be improved or designed differently—for example, by adopting approaches similar to myloasm. Importantly, this constraint reflects the assembly module rather than the MSR sketching technique itself, making it likely addressable through algorithmic adjustments. A promising but vast avenue for enhancing the assembler involves modifying the MSR function, which we designed to be pseudo-random. For instance, we could introduce guarantees based e.g. on syncmers [10] to ensure that at least one base is produced for all windows of length *w*. Another potential improvement could involve exploiting base qualities to estimate and improve the quality of the reduced sequences. The authors of [4] demonstrated that altering the function can significantly enhance results when aligning reads reduced with an MSR of order 2, indicating that the choice of the MSR function has a substantial impact on downstream applications. We hypothesize that a carefully selected MSR function could also enable Alice to effectively handle reads with higher error rates, although the challenge lies in the vast number of MSR functions available for exploration.

While this study concentrates on using MSR sketching for metagenome assembly, the technique has far-reaching potential beyond that scope. Because assembled genomes typically exhibit very low error rates, MSR sketches could be employed for tasks such as indexing or aligning assemblies—e.g., constructing pangenome graphs. Additional promising applications include SNP calling and read alignment, where the efficiency and accuracy of MSR sketching could provide substantial benefits.

## 4 Conclusion

In this study, we present a novel approach for speeding up the assembly of highly precise reads through the introduction of Mapping-friendly Sequence Reductions (MSR) sketches. MSR sketches allow to speed up dramatically genome assembly without sacrificing strain resolution. This method is implemented in a assembler named Alice, which we evaluated on various datasets, including a diploid *Adineta vaga* genome, a challenging mock community comprising five conspecific strains of *Escherichia coli*, a human stool sample and a large sequencing of a soil sample. Alice operated an order of magnitude faster than competing assemblers while maintaining a low memory usage. Moreover, Alice provided the most complete assemblies for high-coverage datasets and is able to distinguish highly similar strains.

## 5 Methods

The fundamental difference between (meta)MDBG and Alice is their sketching scheme. In the next two subsections, we introduce MSR sketches and their interest. In the subsequent sections, we detail the implementation of the Alice assembler.

### 5.1 Mapping-friendly Sequence Reductions (MSR) sketches

Mapping-friendly Sequence Reductions are functions that transform a sequence of characters into a new sequence [4]. A MSR is defined by an alphabet (in this case, the DNA alphabet {*A, C, G, T*}), an order *l* and a transforming function *g* that maps each sequence of length *l*, or *l*-mer, to either a character in the alphabet or a special “empty” character *ϵ*. To ensure that a sequence and its reverse complement are reduced to reverse complement sequences (which is important for genome assembly), an extra constraint is added to *g*: it must map reverse-complement *l*-mers to reverse-complement bases.

MSRs work by taking an input sequence and breaking it down into successive overlapping *l*-mers, which are sequentially passed through the function *g* to produce a reduced sequence. If *g* returns a character, that character is added to the reduced sequence. If *g* returns the empty character *ϵ*, nothing is added to the reduced sequence. The pseudocode for this process is provided in Algorithm 1.

By design, if the length *l* is not too large, two highly similar sequences will share many *l*-mers in the same order, resulting in highly similar reduced sequences. Consequently, the reduced versions of two sequences that align have a high probability of aligning as well; we refer to this property of the reduction as mapping-friendliness. Importantly for us, this mapping-friendly property is also assembly-friendly: the assembly of reduced reads is equivalent to the reduced assembly of the original reads (notwithstanding assembly errors). Reduced reads can thus be used as sketches of their non-reduced counterparts while being potentially much shorter.

#### Algorithm 1

Mapping-friendly Sequence Reductions

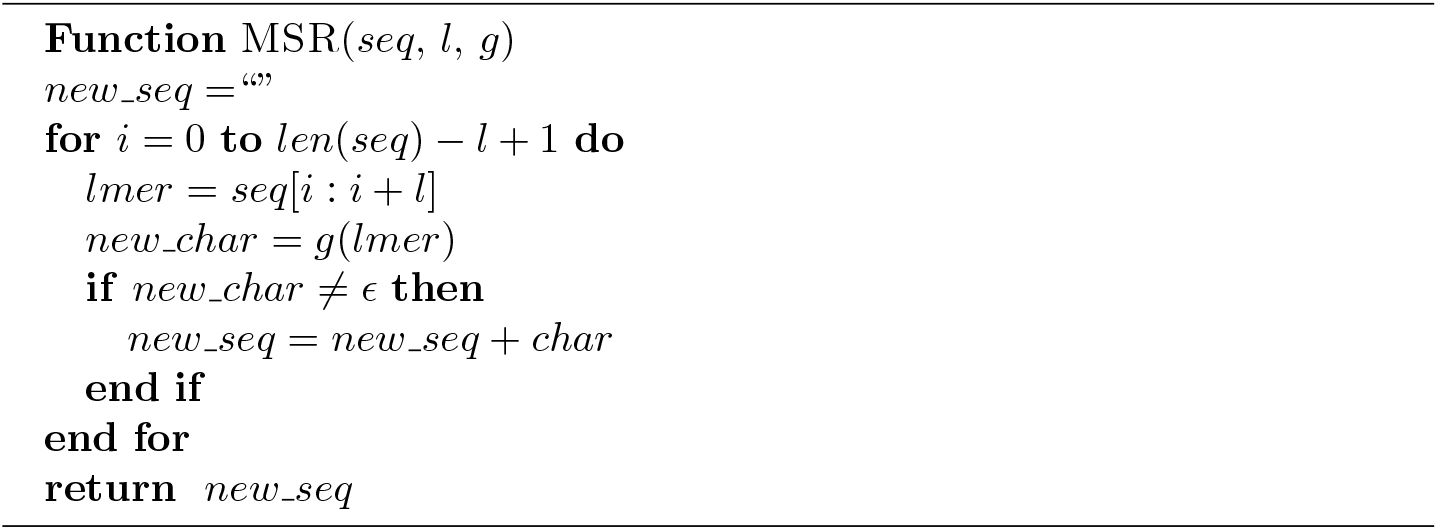

### 5.2 The power of MSR sketches

The main limitation of *k*-mer subsampling techniques such as MinHash [27], FracMinHash [5], or *k*-min-mers [3] is that a change in the input sequence affects the sketch only if it creates or destroys a sampled *k*-mer. By design, at high compression rates, most *k*-mers are not sampled, so most small changes in the input sequences leave the sketch untouched. MSR sketches overcome this limitation by distributing information more evenly. While *k*-mer subsampling retains the full sequence of each selected *k*-mer (i.e., 2*k* bits per selected *k*-mer), an MSR reduces each selected *l*-mer to a single character (2 bits). Consequently, for a given sketch size, an MSR can afford to select far more *l*-mers than a subsampling method selects *k*-mers, and can also afford to use arbitrarily large values of *l*. Together, these two properties allow selected *l*-mers to tile the entire input sequence, ensuring that every base participates in multiple selected *l*-mers. For example, the default settings of *Alice* use *l* = 101 and output 5% of the *l*-mers to non-empty bases, meaning that every base of the input sequence is covered on average by five selected 101-mers. As a result, a single-base change in the input sequence will influence — and likely alter — on average five characters in the output sketch.

As an illustration, let us compare exactly the behavior of MSR sketches and *k*-min-mers showcasing the same compression ratio when confronted with a single base change. We define the compression factor *c* of a sketching method as the expected ratio of the number of bases in a random sequence and the number of bases in its sketch. For MSR sketching, this is equal to the inverse of the ratio of *l*-mers mapping to non-empty characters. For mDBG, following the notation in [11], *c* = 1*/δk*′.

Let us imagine two infinite sequences differing by a single substitution. For the sake of simplicity, let us assume that no *k*′-mer or *l*-mer is repeated around this base. In mBDG, a *k*′-mer has a probability *δ* = 1*/k*′*c* of being sampled, and *k*′ *k*′-mers overlap the SNP. The probability that the *k*-min-mer sketches of the two sequences are different is thus

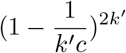

In MSR sketching, two sketches are identical if the *l* consecutive *l*-mers around the SNP on each sequence output the same bases in the same order. The function *g* employed to produce our MSR sketches is crafted to ensure that there is virtually no correlation between input *k*-mers and their corresponding image through *g* (see Methods). Therefore we can compute the probability of the two sketches being identical as the probability that they have the same number of bases *i* (which is given by the square of the probability of choosing *i* items among *l*, if each of them has a probability 1*/c* of being chosen) multiplied by the probability that two series of *i* DNA bases are identical (which is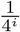):

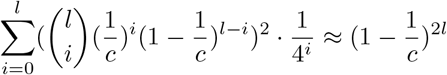

If we use Alice’s default compression factor of 20 and order *l* of 101, the probability that the mDBG sketches of the two sequences are different is of less than 10%, whereas the probability that the MSR sketches of the two sequences are different is higher than 99%.

The assembly process is illustrated Figure 1.

### 5.3 Reducing input reads

All reads are initially reduced using a Mapping-friendly Sequence Reduction (MSR) provided by Alice. The MSR allows the user to select the order *l* (default value of 101) and the compression factor *c* (default value of 20). The *l*-mers of the reads are processed through a function *g*. This function takes an *l*-mer as input and outputs either a single base, which is appended to the growing reduced read, or an “empty” base *ϵ*, which is not appended to the growing reduced read.

The function *g* of the MSR is designed as follows. The *l*-mer is converted into its canonical form, which is either the original *l*-mer or its reverse complement if the reverse complement is lexicographically smaller. *g* then applies to the canonical *l*-mer a pseudo-random hash function yielding a hash between 0 and 1 [15]. It distinguishes five cases:

- if the hash is smaller than 1/2*c* and the original *l*-mer is canonical, an *A* is outputted;
- if the hash is smaller than 1/2*c* and the original *l*-mer is not canonical, a *T* is outputted;
- if the hash is between 1/2*c* and 1/*c* and the original *l*-mer is canonical, a *C* is outputted;
- if the hash is between 1/2*c* and 1/*c* and the original *l*-mer is not canonical, a *G* is outputted;
- if the hash is between 1/*c* and 1, *ϵ* is outputted.

This MSR is combined with a classic homopolymer compression process that occurs before all the reads are sketched, at the very beginning of the process, to reduce the error rate of the reads.

### 5.4 Assembling reduced reads

Several existing assemblers were evaluated for assembling reduced reads. Long-read assemblers generally struggled with the short reduced reads, while short-read assemblers performed considerably better. Among them, metaSPAdes [26] produced high-quality assemblies; however, it collapsed closely related strains to enhance continuity—a behavior that undermines the strain-resolution objective of Alice. Although metaSPAdes can be used as an optional backend in Aliceasm, all results presented here are based on a custom assembler we developed. Our custom assembler operates in three steps.

1. **Read correction**. We follow a well-established procedure used in, e.g., [20, 1]: a De Bruijn Graph (DBG) is first constructed from the reads using BCALM2 [8]. Sequencing errors are then removed from the graph by discarding all *k*-mers observed only once, as well as tips and bubbles composed exclusively of *k*-mers observed fewer than 5 times—a standard technique employed in, e.g., [11, 20, 2]. When multiple low-coverage bubbles occur within a window of 10 *k* bases of one another, they are removed only if `--single-genome` mode is activated, as they may otherwise represent rare haplotypes in metagenomic settings. Finally, the original reads are aligned back onto the error-corrected graph, and each read sequence is replaced by the sequence of the contig(s) it aligns to.
2. **Assembly**. The corrected reads are assembled using De Bruijn graphs of increasing *k*-mer sizes, a classical iterative strategy implemented in, e.g., [26].
3. **Graph untangling**. The final De Bruijn graph is untangled to improve continuity and to duplicate unitigs that appear multiple times in the genome, following the approach of Unicycler [38]. Specifically, all corrected reads are first aligned onto the assembly graph. A contig is then classified as a *single-copy contig* if all reads aligning to it do so consensually, forming a unique path on both its left and right flanks. When two single-copy contigs are consistently linked by a set of reads, the contigs along the connecting path are duplicated, merging the two single-copy contigs into a single, longer, single-copy contig.

### 5.5 Recovering the uncompressed assembly

The compressed assembly represents the reduced version of the final assembly. Inflating this reduced version back to the full assembly is not straightforward, as the MSR reduction function is not invertible.

Our method involves three steps:

- creating an inventory of *k*-mers that tile the compressed assembly, using a *k*-mer size of 31 by default. For example, two 3-mers that tile the sequence “ACCGTT” are “ACC” and “GTT”;
- re-running the MSR on all original reads, and each time a tiling *k*-mer is produced, record the corresponding uncompressed sequence. For example, we can record that “ACC” corresponds to “GTCGCATGACTGAT” and “GTT” to “TCCGACTCATCAGA”; and finally
- reconstructing the full assembly by concatenating the uncompressed sequences of the tiling *k*-mers, which would yield in our example “GTCGCATGACTGATCCGACTCATCAGA”.

### 5.6 Choice of parameters

We experimented with different parameter choices for the compression factor and the order of reduction on the ZymoBIOMICS Gut Microbiome Standard dataset to understand how these parameters influence the final assembly.

We conducted two experiments: one to assess the effects of the order and another to assess the effect of the compression factor. First, we tested compression factors of 100, 50, 20, 10, and 5 with an order of 101. Second, we tested orders of 11, 21, 51, 101, and 201 with a compression factor of 10.

The variation of these parameters primarily impacted the completeness of the resulting assemblies and the run-times of the pipelines, while their accuracy, duplication ratio, and continuity remained equivalent.

As expected, the run-time increased with the compression factor, as there was more data to assemble. This is illustrated in Figure 5.

**Figure 5:**
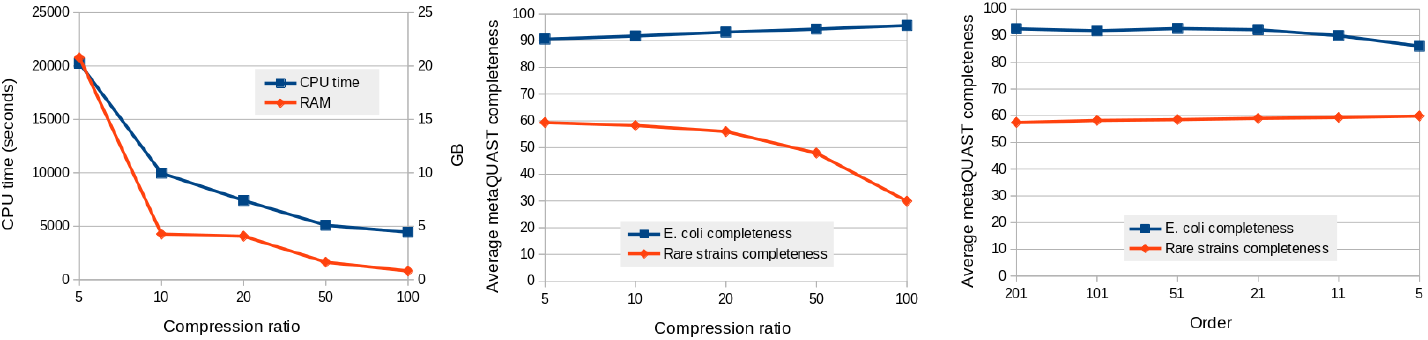
Variation of resource usage and metaQUAST completeness of the ZymoBIOMICS gut microbiome standard assemblies with different compression factor and orders. “Rare strains” refer to *C. albicans, S. cerevisiae*, and *S. enterica*. The completeness displayed are arithmetic means of the completeness of the different genomes of the categories.

Compressing more the data also had a positive impact on the completeness and continuity of the five highly similar *E. coli* strains (Figure 5). When investigating the 27-mer completeness (not shown), all assemblies had a similar amount of missing 27-mers. Hence, the main difference explaining the difference in completeness was that repeated regions were more shrunk when the data were more compressed, which helped to assemble repeated regions closer to their true multiplicity and thereby improved continuity.

However, the results represented in Figure 5 show that compression negatively impacted the completeness of the *C. albicans, S. cerevisiae*, and *S. enterica* genomes, which had relatively low coverage (see Supplementary Table 1 for the coverages). This is because the assembly algorithm requires a sufficient number of error-free 25-mers in the compressed reads to effectively correct reads and produce a complete assembly. An error-free compressed 25-mer corresponds to an error-free uncompressed sequence of average length 25 \**c* + *l* without errors. Therefore, as the compression factor *c* increases, the number of correct 25-mers in the reads decreases. For genomes with low coverage, high compression can result in the loss of precious 25-mers, leading to insufficient coverage of some regions, hindering their assembly.

The order was found to have relatively little impact on the resulting assemblies. The only significant effect observed was when the order decreased to 11 and 5, where *l/c* approached or fell below 1. In these cases, the assembler began collapsing highly similar sequences, leading to a decrease in the completeness of the five *E. coli* strains. Despite the fact that the error rate scales approximately linearly with *l*, increasing *l* did not have a significant negative impact on the completeness of the assemblies. This is because the errors in the compressed reads cluster in increasingly large clusters, but the error-free regions between these clusters diminish in size only slowly with *l*.

## Supporting information

in the supplementary data set

## 6 Declarations

## Acknowledgments

We acknowledge the GenOuest bioinformatics core facility (https://www.genouest.org) for providing the computing infrastructure. The programs Tablet [24] and Bandage [37] were used to visualize data while developing Alice. The DNA extracts used for sequencing were kindly provided by Karine Van Doninck and Julie Virgo.

We thank Rayan Chikhi for his proofreading and advice.

For the purpose of open access, the authors have applied a CC-BY public copyright license to any Author Manuscript version arising from this submission.

## Availability of data and materials

Alice is freely available on github at https://www.github.com/RolandFaure/aliceasm(and on Zenodo with the DOI 10.5281/zenodo.19737094). Alice is written in C++ and is distributed under the Affero-GPL3.

All the datasets used for benchmarking Alice are available publicly, under accession numbers SRR13128013 for the ZymoBIOMICS Gut Microbiome Standard, SRR28996637 for the human gut microbiome dataset, ERR15289804 for the soil and BioProject PRJNA1335825 for *Adineta vaga*. Zymo-HiFi mock reference genomes are available at https://s3.amazonaws.com/zymofiles/BioPool/D6331.refseq.zip

Assemblies were run with Alice-asm version 0.7.14, hifiasm 0.24.0-r702, Flye 2.9.5-b1801, metaMDBG 1.0, LJA commit 99f93262c. All assemblies and command lines used are available in Zenodo, DOI 10.5281/zenodo.17179435.

For the purpose of Open Access, a CC-BY public copyright licence has been applied by the authors to the present document and will be applied to all sub-sequent versions up to the Author Accepted Manuscript arising from this sub-mission.

## Competing interests

None declared.

## Funding

RF is supported by the Horizon Europe ERC grant number 101088572 “IndexThePlanet”. HiFi sequencing of Adineta vaga was funded by the Horizon 2020 research and innovation program of the European Union under the Marie Sklodowska-Curie grant agreement No 764840 (ITN IGNITE, www.itnignite.eu) to JFF.

## Author’s contributions

**Roland Faure:** Investigation, Conceptualization, Software, Writing. **Baptiste Hilaire:** Investigation. **Dominique Lavenier:** Supervision. **Jean-Françcois Flot:** Supervision.

## Notes

https://doi.org/10.5281/zenodo.17179435

### Competing Interest Statement

The authors have declared no competing interest.

### Summary of Updates

The software has been improve, the results have been changed.

