## Supplementary Tables 1 and 2 for "Fast and haplotype-aware assembly of high-fidelity reads based on MSR sketching: the Alice assembler"

April 27, 2026

### A Completeness and duplication ratio of the Zymobiomics Gut Standard assemblies

| Genomes | coverage | Alice | hifiasm_meta | metaFlye | metaMDBG | myloasm |
|---|---|---|---|---|---|---|
| *V. rogosae* | 1667 | 1.0 / 1.0 | 1.0 / **49** | 1.0 / 1.0 | 1.0 / 1.0 | 1.0 / 1.0 |
| *F. prausnitzii* | 1250 | 1.0 / 1.0 | 1.0 / **28** | 1.0 / 1.0 | 1.0 / 1.0 | 1.0 / 1.0 |
| *R. hominis* | 1206 | 1.0 / 1.0 | 1.0 / **2.3** | 1.0 / 1.0 | 1.0 / 1.0 | 1.0 / 1.0 |
| *L. fermentum* | 789 | 1.0 / 1.0 | 1.0 / **1.4** | 1.0 / 1.0 | 1.0 / 1.0 | 1.0 / 1.0 |
| *B. adolescentis* | 750 | 1.0 / 1.2 | 1.0 / **2.5** | 1.0 / 1.0 | 1.0 / 1.0 | 1.0 / 1.0 |
| *B. fragilis* | 700 | 1.0 / 1.1 | 1.0 / **15** | 1.0 / 1.0 | 1.0 / 1.0 | 1.0 / 1.0 |
| *F. nucleatum* | 625 | 1.0 / 1.0 | 1.0 / **5.9** | 1.0 / 1.0 | 1.0 / 1.0 | 1.0 / 1.0 |
| *P. corporis* | 517 | 0.99 / 1.0 | 1.0 / **10** | 1.0 / 1.0 | 1.0 / 1.0 | 1.0 / 1.0 |
| *E. coli* JM109 | 152 | 1.0 / 1.1 | 1.0 / **2.1** | **0.44** / 1.1 | **0.38** / 1.0 | **0.62** / 1.0 |
| *E. coli* B3008 | 152 | 1.0 / 1.0 | 1.0 / **1.6** | **0.83** / 1.0 | **0.36** / 1.0 | 1.0 / 1.0 |
| *E. coli* B1109 | 148 | 1.0 / 1.2 | 0.99 / **2.1** | **0.60** / 1.1 | **0.78** / 1.0 | 0.99 / 1.0 |
| *E. coli* b2207 | 140 | 0.97 / **1.9** | 0.99 / **2.2** | **0.47** / 1.0 | **0.37** / 1.0 | 1.0 / 1.0 |
| *E. coli* B766 | 140 | 0.96 / 1.0 | 0.96 / **1.5** | 0.95 / 1.0 | 0.96 / 1.0 | 0.96 / 1.0 |
| *A. muciniphila* | 133 | 1.0 / 1.0 | 1.0 / **1.6** | 1.0 / 1.0 | 1.0 / 1.0 | 1.0 / 1.0 |
| *C. difficile* | 91 | 1.0 / 1.0 | 0.91 / **1.3** | 1.0 / 1.0 | 1.0 / 1.0 | 1.0 / 1.0 |
| *C. albicans* | 28 | **0.60** / 1.2 | **0.68** / 1.2 | **0.72** / 1.0 | **0.81** / 1.0 | **0.79** / 1.0 |
| *S. cerevisiae* | 27 | **0.58** / 1.2 | **0.73** / 1.2 | **0.80** / 1.0 | **0.89** / 1.0 | **0.80** / 1.0 |
| *M. smithii* | 14 | 1.0 / 1.0 | 1.0 / 1.0 | 1.0 / 1.0 | 1.0 / 1.0 | 1.0 / 1.0 |
| *S. enterica* | 1 | **0.07** / 1.0 | **0.12** / 1.0 | **0.02** / 1 | **0.24** / 1.0 | **0.10** / 1.0 |
| *C. perfringens* | 1 | **0** / - | **0** / - | **0** / - | **0** / - | **0** / - |
| *E. faecalis* | 0 | **0** / - | **0** / - | **0** / - | **0** / - | **0** / - |

Table 1: Completeness / duplication ratios of the alice_07, hifiasm_meta, metaFlye, metaMDBG and myloasm assemblies with respect to all the genomes of the mix, ordered by decreasing coverage and computed by metaQUAST. Completeness values below 0.9 and duplication values above 1.2 correspond to perfectible assemblies and are highlighted in red. Note that *C. albicans* and *S. Cerevisiae* are yeast and all other species are bacteria.

### B Contiguities

| Genomes | coverage | Alice | metaFlye | hifiasm_meta | metaMDBG | myloasm |
|---|---|---|---|---|---|---|
| *V. rogosae* | 1667 | 1.38 | 1.84 | 0.03 | 1.85 | **2.04** |
| *F. prausnitzii* | 1250 | 2.05 | 1.91 | 0.02 | 2.00 | **2.72** |
| *R. hominis* | 1206 | **3.10** | 2.82 | 2.30 | 2.70 | 2.56 |
| *L. fermentum* | 789 | 1.73 | 1.72 | 1.61 | **1.74** | **1.74** |
| *B. adolescentis* | 750 | **1.56** | **1.56** | 0.63 | 1.46 | **1.56** |
| *B. fragilis* | 700 | 0.95 | **4.79** | 0.53 | 1.94 | 2.94 |
| *F. nucleatum* | 625 | 1.86 | **2.15** | 2.01 | 1.26 | 2.14 |
| *P. corporis* | 517 | 0.60 | 1.06 | 2.10 | 1.58 | **2.44** |
| *E. coli* JM109 | 152 | 0.32 | 0.02 | 0.14 | - | **0.54** |
| *E. coli* B3008 | 152 | 1.63 | 0.06 | 0.96 | - | **3.78** |
| *E. coli* B1109 | 148 | 0.47 | 0.01 | 0.13 | 0.46 | **1.22** |
| *E. coli* b2207 | 140 | 0.09 | - | 0.22 | - | **3.98** |
| *E. coli* B766 | 140 | 0.33 | 0.22 | 0.36 | 0.36 | **0.39** |
| *A. muciniphila* | 133 | **2.78** | 2.45 | 2.22 | 1.56 | 2.05 |
| *C. difficile* | 91 | 2.54 | 3.84 | 3.18 | 2.29 | **3.99** |
| *C. albicans* | 28 | 0.01 | 0.03 | 0.05 | 0.06 | **0.07** |
| *S. cerevisiae* | 27 | 0.02 | 0.04 | 0.07 | **0.12** | 0.11 |
| *M. smithii* | 14 | 0.23 | 0.51 | 0.41 | 1.20 | **1.15** |
| *S. enterica* | 1 | - | - | - | - | - |
| *C. perfringens* | 1 | - | - | - | - | - |
| *E. faecalis* | 0 | - | - | - | - | - |

Table 2: NGA50 of the genomes of the Zymobiomics gut standard reconstructed by the different assemblers, measured with the metaQUAST. All the figures are in Mbp. Empty cells correspond to genomes that were not at least 50% covered. The best values are highlighted in bold.
